# *Xanthomonas* type III effector XopN targets scaffold protein OsRACK1B to suppress rice immunity

**DOI:** 10.1101/2025.11.03.686189

**Authors:** Jiuxiang Wang, Zhe Ni, Xin Chen, Yaqi Zhang, Huajun Qin, Haoyu Wang, Yongqiang He, Jiliang Tang, Sheng Huang

**Author notes:** These authors have contributed equally to this work (Jiuxiang Wang, Zhe Ni).

## Abstract

Bacterial leaf streak (BLS), caused by *Xanthomonas oryzae* pv. *oryzicola* (*Xoc*), is a major bacterial disease that poses a substantial threat to rice production. Although *Xoc* deploys a variety of effectors to subvert host defenses, the mechanistic basis of how these effectors modulate plant immune signaling remains largely unclear. Here, we show that XopN suppresses rice immune responses and physically interacts with the rice scaffold protein OsRACK1B. Notably, *OsRACK1B* expression is induced following *Xoc* infection in a type III secretion system (T3SS)-independent manner. Functional analyses indicate that *OsRACK1B* overexpression enhances *Xoc* resistance, whereas loss-of-function mutants exhibit heightened susceptibility. This increased susceptibility is characterized by compromised immunity, including reduced callose deposition, diminished reactive oxygen species (ROS) accumulation, and attenuated expression of defense-related genes such as *OsPBZ1* and *OsPAL1*. Furthermore, we demonstrate that XopN competes with OsRap2.6 for binding to OsRACK1B, thereby disrupting the formation of the OsRACK1B–OsRap2.6 immune complex. These findings reveal a novel virulence strategy in *Xoc* and identify promising gene targets for developing disease-resistant rice varieties.

## Introduction

Rice (*Oryza sativa* L.) is a staple crop that sustains the nutritional demands of more than half of the global population (Yuan et al., 2021). However, its productivity is continually threatened by a diverse array of pathogens and pests, which compromise both yield and grain quality (He et al., 2023). Among these, bacterial leaf streak (BLS), caused by *Xanthomonas oryzae* pv. *oryzicola* (*Xoc*), is a major bacterial disease that results in yield losses ranging from 8% to 32% (Liu et al., 2014). Genetically encoded resistance remains the most effective and environmentally sustainable approach for long-term disease management. Despite progress, only a limited number of resistance (*R*) genes, such as *Xo1*, *xa5*, *bls1*, and *OsBLS6.2*, have been identified as effective against BLS. (Ma et al., 2021; Triplett et al., 2016; Xie et al., 2023; Xie et al., 2014). This underscores the pressing need to elucidate the molecular strategies employed by *Xoc* to facilitate infection, which will be instrumental for the development of novel resistance traits.

The continuous co-evolution of pathogenic microbes and their host plants has led to the emergence of a sophisticated, two-layered innate immune system in plants: pattern-triggered immunity (PTI) and effector-triggered immunity (ETI) (Jones and Dangl, 2006; Wang et al., 2022). PTI is initiated upon recognition of conserved pathogen-associated molecular patterns (PAMPs) by pattern recognition receptors (PRRs) located at the plasma membrane (Li et al., 2016). In contrast, ETI is activated intracellularly through nucleotide-binding leucine-rich repeat (NLR) receptors that detect pathogen-secreted effectors (Tsuda and Katagiri, 2010). Immune activation induces a range of defense responses, including the production of reactive oxygen species (ROS), callose deposition, mitogen-activated protein kinase (MAPK) cascade activation, upregulation of defense-related genes, and hypersensitive cell death (HR) at the site of infection (Gao et al., 2021; Spoel and Dong, 2012; Wang et al., 2020). While PTI provides basal immunity, ETI elicits a more robust and rapid defense, often accompanied by localized cell death to restrict pathogen spread (Spoel and Dong, 2012).

Many bacterial phytopathogens utilize a type III secretion system (T3SS) to translocate type III-secreted effectors (T3SEs) directly into host cells, where they manipulate host immunity and facilitate colonization (Boller and He, 2009). In *Xanthomonas* species, T3SEs are broadly classified into transcription activator-like (TAL) effectors and non-TAL effectors (Kay and Bonas, 2009). TAL effectors function as transcriptional activators upon nuclear import, often reprogramming host gene expression to support bacterial growth. For instance, TAL effectors from *X. oryzae* pv. *oryzae* (*Xoo*) induce *OsSWEET* gene expression, enabling pathogen exploitation of host sugars (Oliva et al., 2019). Similarly, the TAL effector Tal2g from *Xoc* promotes disease by upregulating the sulfate transporter gene *OsSULTR3;6* (Cernadas et al., 2014).

By contrast, non-TAL effectors exploit a diverse array of biochemical strategies to reprogram host cellular processes and facilitate bacterial colonization. For example, *X. campestris* pv. *vesicatoria* (*Xcv*) effector XopL functions as an E3 ubiquitin ligase that interacts with and degrades the autophagy component SH3P2, thereby promoting infection (Leong et al., 2022). Another effector, XopR from *X. campestris* pv. *campestris* (*Xcc*), forms biomolecular condensates via electrostatic-driven liquid–liquid phase separation (LLPS), coacervating with formin-mediated actin nucleation complexes and manipulating the host cytoskeleton (Sun et al., 2021). Recent studies have increasingly highlighted the functional diversity of non-TAL effectors deployed by *Xoc*. For instance, XopC2 acts as a kinase that phosphorylates the Skp1 protein OSK1, thereby suppressing immune signaling (Wang et al., 2021). The effector AvrRxo1 degrades the pyridoxal phosphate synthase OsPDX1, reducing vitamin B6 levels and suppressing stomatal immunity (Liu et al., 2022). XopAP competitively binds phosphatidylinositol 3,5-bisphosphate (PtdIns(3,5)P2), inhibiting vacuolar acidification by interfering with vacuolar H⁺-pyrophosphatase (V-PPase), ultimately promoting stomatal opening(Liu et al., 2022). More recently, XopAK was shown to repress *OsMYBxoc1* transcription and alter host iron homeostasis, thereby modulating rice immunity (Zhang et al., 2024).

Several non-TAL effectors, including AvrBs2, hrpE3 and XopN, have been identified as critical for the full virulence of *Xoc* (Cui et al., 2013; Li et al., 2015; Liao et al., 2020). However, the molecular mechanisms by which these effectors contribute to pathogenicity in rice remain largely unknown. Our previous study established that XopN significantly enhances the virulence of the *Xoc* strain GX01 (Liao et al., 2020). Here, we demonstrate that XopN suppresses rice immune responses and identify the rice scaffold protein OsRACK1B as a novel host target of XopN. Mechanistically, XopN impairs OsRACK1B function by disrupting its association with the transcription factor OsRap2.6, thereby dismantling a critical immune signaling complex. Functional analyses revealed that overexpression of *OsRACK1B* enhances rice resistance to *Xoc*, whereas *OsRACK1B* knockout lines exhibit heightened susceptibility and impaired immune responses. Collectively, these findings highlight a virulence strategy whereby XopN targets a central immune scaffold to suppress host defense, highlighting OsRACK1B as a promising molecular target for breeding BLS-resistant rice varieties.

## Results

### Δ*xopN* mutant elicits elevated rice immune responses

To gain insights into the evolutionary conservation of XopN, we performed a BLAST analysis using its amino acid sequence as a query. The results revealed that XopN is highly conserved within the *Xanthomonas* genus, exhibiting 96.0% sequence identity with its homolog in *Xoo*, 81.7% identity with the counterpart in *Xcc*, and 60.9% identity with the homolog in *Xcv* (Figure S1). Notably, a putative homolog sharing 52.5% identity was also identified in *Acidovorax citrulli*, a vascular pathogen of cucurbits, suggesting potential horizontal conservation beyond the *Xanthomonas* clade. In contrast, XopN shows limited sequence similarity to effector-like proteins from more distantly related phytopathogens, such as *Ralstonia solanacearum* (25.2% identity), indicating that XopN and its functional equivalents are largely restricted to *Xanthomonas* and a few phylogenetically proximate taxa.

We further investigated the functional role of the *Xoc* effector XopN in modulating rice innate immunity by comparing host responses triggered by the wild-type strain GX01, the Δ*xopN* mutant, and the complemented strain CΔ*xopN*. Callose, a β-1,3-glucan polymer, serves as a critical component of plant immune defenses by reinforcing cell walls to restrict pathogen ingress and dissemination (Li et al., 2023). Accordingly, we first assessed callose deposition in rice leaves infiltrated with GX01, Δ*xopN*, or CΔ*xopN*. The results revealed that leaves infected with the Δ*xopN* mutant exhibited significantly enhanced callose accumulation relative to those inoculated with either GX01 or CΔ*xopN* (Figure 1a,b). We next assessed ROS production, a rapid defense response typically associated with PTI activation. Using 3,3′-diaminobenzidine (DAB) staining, we observed a marked increase in H₂O₂ accumulation in Δ*xopN*-infected leaves compared to those infected with GX01 or CΔ*xopN* (Figure 1c). To determine whether XopN also modulates transcriptional defense responses, we conducted quantitative reverse transcription PCR (RT–qPCR) to analyze the transcript levels of canonical defense marker genes. The expression of *OsPBZ1* and *OsPAL1* was significantly upregulated in Δ*xopN*-infected leaves compared to GX01 (Figure 1d,e), indicating that XopN suppresses transcriptional activation of these defense-associated genes.

**Figure 1.**
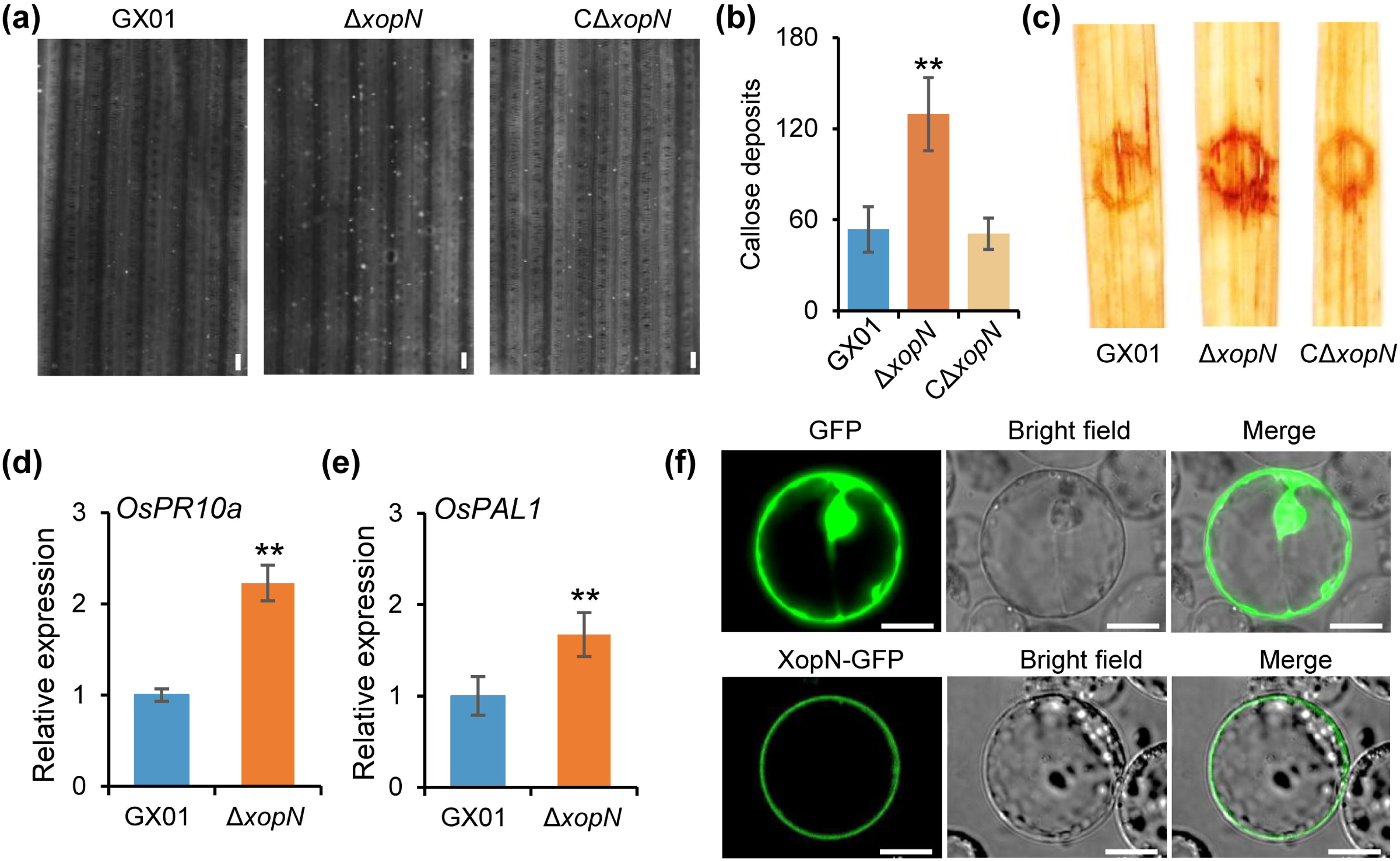
XopN suppresses immune responses in rice. **(a)** Callose deposition in rice leaves at 72 hours post-inoculation with the *Xoc* wild-type strain GX01, the Δ*xopN* mutant, and the complemented strain CΔ*xopN*. Scale bar, 100 μm. **(b)** Quantification of callose deposits per leaf. Data represent means ± standard deviations, n=5. Asterisks denote statistically significant differences between the Δ*xopN* mutant and either the wild-type or complemented strain (**, *P* < 0.01, Student’s *t*-test). **(c)** DAB staining of rice leaves at 72 hours post-inoculation with GX01, Δ*xopN*, and CΔ*xopN*. **(d, e)** Relative expression levels of *OsPBZ1* and *OsPAL1* in NIP rice at 24 hours post-inoculation with GX01, Δ*xopN*, and CΔ*xopN*. Rice actin was used as an internal control. Data represent means ± standard deviations, n = 3. Asterisks indicate significant differences compared to wild-type or complemented strains (**, *P* < 0.01, Student’s *t*-test). **(f)** Subcellular localization of XopN in planta. Rice protoplasts were transfected with a construct expressing XopN fused to the N-terminus of GFP (XopN-GFP). At 24 h post-transfection, fluorescence was visualized by confocal laser scanning microscopy. Scale bars, 10 μm.

Effectors are translocated into host cells to target various subcellular compartments, manipulating diverse physiological and immune processes (Lo Presti et al., 2015). To determine the subcellular localization of XopN, we expressed XopN-GFP in rice protoplasts. The fusion protein was primarily localized to the plasma membrane (PM) and cytoplasm (Figure 1f). Together, these results demonstrate that XopN functions to dampen rice immune responses at multiple levels, including callose deposition, ROS production, and defense gene induction.

### XopN targets the scaffold protein OsRACK1B in rice

To identify potential rice targets of XopN, we conducted a yeast two-hybrid (Y2H) screen using a rice cDNA library with XopN as bait. This screen identified OsRACK1B, a scaffold protein, as a putative interactor of XopN. To confirm this preliminary interaction, full-length coding sequences of XopN and OsRACK1B were cloned into the pGBKT7 (bait) and pGADT7 (prey) vectors, respectively. Y2H assays confirmed that XopN interacts with OsRACK1B in yeast (Figure 2a). To further substantiate this interaction, we performed a reciprocal Y2H assay in which full-length OsRACK1B was cloned into the pGBKT7 vector and XopN into pGADT7. This reversed configuration also yielded a positive interaction signal (Figure 2a). To assess whether this interaction occurs in planta, we performed a bimolecular fluorescence complementation (BiFC) assay in rice protoplasts. Yellow fluorescence was observed only upon co-expression of nYFP–XopN and RACK1B–cYFP (Figure 2b), indicating a specific interaction between the two proteins. Additionally, a pull-down assay was conducted to validate the interaction in vitro. His-tagged XopN was specifically pulled down by GST–RACK1B, but not by GST alone, further supporting the direct interaction (Figure 2c).

**Figure 2.**
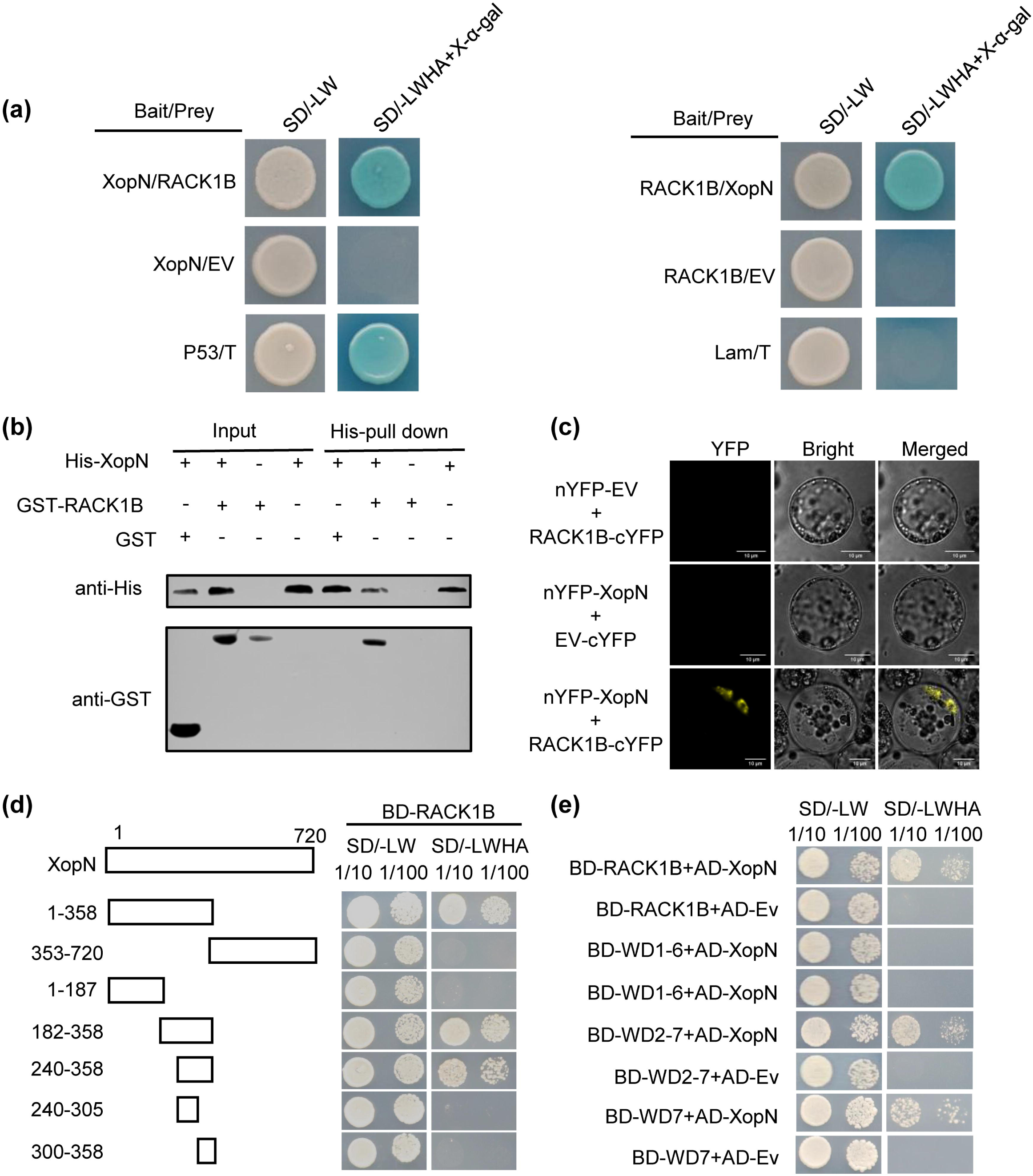
XopN physically interacts with OsRACK1B. **(a)** Y2H analysis showing the interaction between XopN and OsRACK1B. Yeast strain Y2HGold was co-transformed with the indicated plasmid combinations and cultured on SD/-Leu-Trp and the selective medium SD/-Leu-Trp-His-Ade. Plates were incubated at 30 °C for 3 days prior to imaging. The p53/T combination served as a positive control, and Lam/T as a negative control. EV, empty vector control. SD/-LW, SD/-Leu-Trp; SD/-LWHA, SD/-Leu-Trp-His-Ade. **(b)** BiFC assay demonstrating the in vivo interaction of XopN with OsRACK1B in rice protoplasts. XopN was fused to the N-terminal fragment of YFP (nYFP), and OsRACK1B to the C-terminal fragment (cYFP). Reconstituted YFP fluorescence was visualized by confocal microscopy 20 hours post-transformation. Scale bar, 10 μm. EV, empty vector control. **(c)** In vitro pull-down assay confirming the physical interaction between XopN and OsRACK1B. Recombinant His-tagged XopN and GST-tagged OsRACK1B, expressed and purified from *E. coli*, were subjected to pull-down using Ni-NTA resin. Bound proteins were detected by immunoblotting with anti-GST and anti-His antibodies. **(d)** Y2H assay showing the interaction between OsRACK1B and truncated fragments of XopN. **(e)** Y2H assay demonstrating the interaction between XopN and truncated variants of OsRACK1B. Yeast strain Y2HGold was co-transformed with the indicated plasmid combinations and cultured on SD/-Leu-Trp and the selective medium SD/-Leu-Trp-His-Ade. Plates were incubated at 30 °C for 3 days prior to imaging. SD/-LW, SD/-Leu-Trp; SD/-LWHA, SD/-Leu-Trp-His-Ade.

We next mapped the interaction domains between XopN and OsRACK1B using the Y2H system. A series of XopN truncations fused to GAL4-AD were co-expressed with BD-OsRACK1B in yeast. The results indicated that the C-terminal region of XopN (amino acids 240–358) is necessary for the interaction (Figure 2d). Similarly, truncated OsRACK1B constructs were tested for their ability to bind XopN, revealing that the WD7 domain alone was sufficient to mediate the interaction (Figure 2e). Together, these data demonstrate that XopN physically interacts with OsRACK1B both in vivo and in vitro and that this interaction is mediated by specific regions within each protein.

### *OsRACK1B* gene expression is upregulated upon *Xoc* infection

OsRACK1B encodes a 336-amino-acid protein containing seven WD40 repeats, a domain architecture highly conserved across eukaryotic organisms (Figure S2a). Phylogenetic analysis revealed that OsRACK1B shares the highest sequence identity with its paralog OsRACK1A in rice (83.8%) and displays greater similarity to orthologs in monocot species, including maize ZmRACK1 (82.5%) and wheat TaRACK1A (80.5%), than to those in dicots such as tobacco NtRACK1 (68.8%) and *Arabidopsis* AtRACK1C (68.4%) (Supplementary Figure S4B). Moderate sequence conservation was also observed with human (61.7%) and yeast (44.0%) RACK1 homologs (Figure S2b), underscoring a deep evolutionary relationship within the eukaryotic lineage. Notably, no RACK1 homologs have been identified in bacterial genomes, suggesting that RACK1 family proteins are unique to eukaryotes.

To determine whether *Xoc* infection alters *OsRACK1B* transcript levels, we analyzed *OsRACK1B* mRNA abundance in rice leaves following inoculation with the wild-type *Xoc* strain GX01. RT–qPCR revealed that *OsRACK1B* expression was significantly induced at 12 and 24 hours post-inoculation (hpi) (Figure 3a), indicating transcriptional upregulation in response to pathogen challenge. To examine whether this induction depends on the T3SS, we examined *OsRACK1B* mRNA levels in rice leaves inoculated with a T3SS-deficient mutant (Δ*hrcC*). The expression of *OsRACK1B* was induced to a similar level compared to the leaves infected with GX01 (Figure 3b), suggesting that the induction of *OsRACK1B* is independent of the bacterial T3SS.

**Figure 3.**
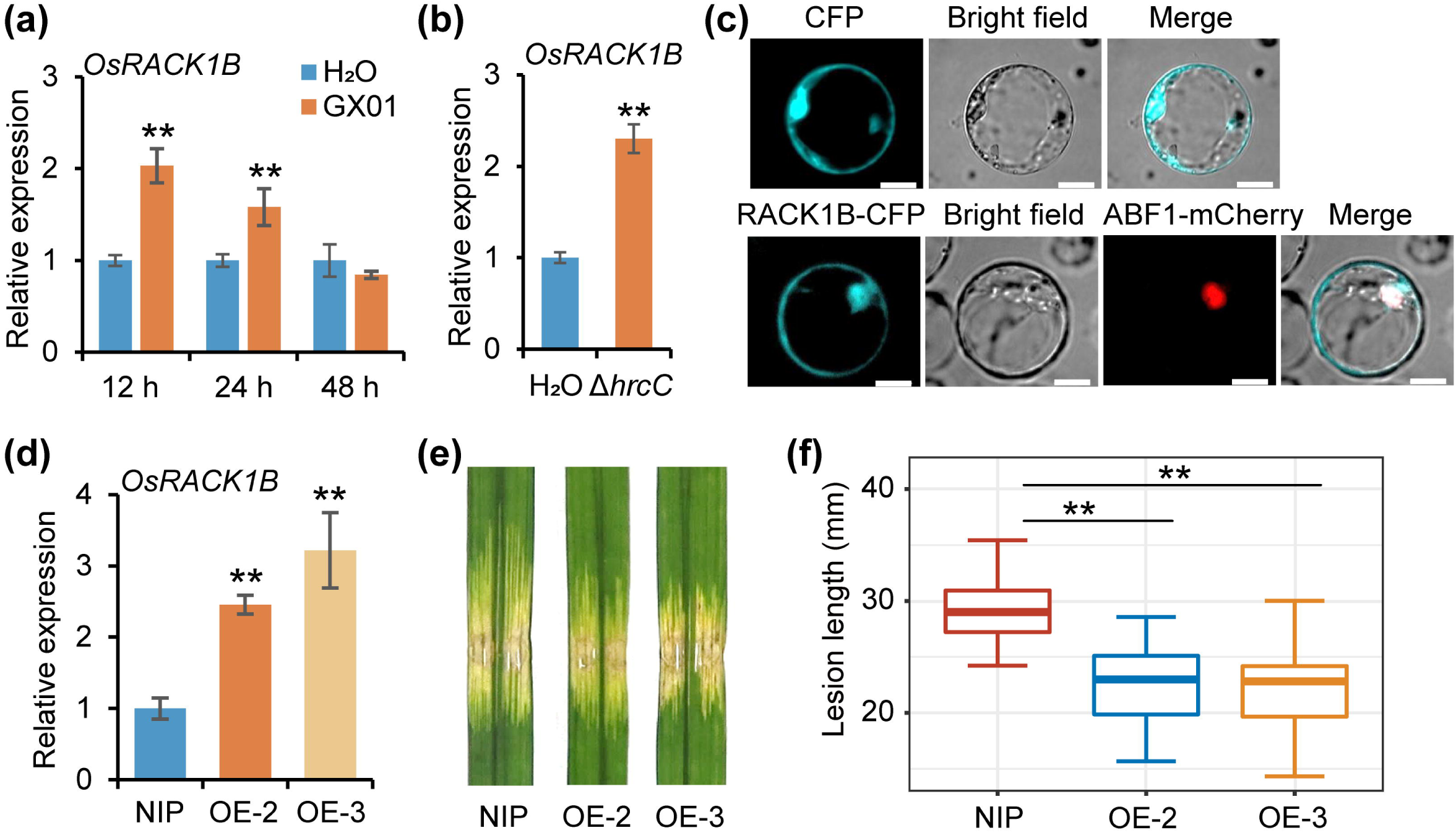
Overexpression of *OsRACK1B* enhances rice resistance to *Xoc*. **(a, b)** Expression dynamics of *OsRACK1B* in rice leaves following inoculation with the wild-type *Xoc* strain GX01 or the T3SS-deficient mutant strain Δ*hrcC*. Rice actin was used as an internal control. Data represent means ± standard deviations, n = 3. Asterisks indicate statistically significant differences (**, *P* < 0.01, Student’s *t*-test). **(c)** Subcellular localization of OsRACK1B in planta. Rice protoplasts were transfected with a construct expressing OsRACK1B fused to the N-terminus of CFP (OsRACK1B-CFP) and co-expressed with the nuclear marker ABF1-mCherry. Fluorescence was detected by confocal laser scanning microscopy at 24 h post-transfection. Scale bars, 10 μm. **(d)** Relative expression of *OsRACK1B* in wild-type NIP and transgenic overexpression lines. Rice actin was used as an internal control. Data represent means ± standard deviations, n = 3. Asterisks indicate statistically significant differences (**, *P* < 0.01, Student’s *t*-test). **(e)** Phenotypes of lesion expansion in the NIP and *OsRACK1B* overexpression lines at 14 days post-inoculation with GX01. **(f)** Lesion lengths in the NIP and *OsRACK1B* overexpression lines at 14 days post-inoculation with GX01. Data are displayed as boxplots with individual data points. Error bars represent maximum and minimum values. Center line, median; box limits, 25th and 75th percentiles. Asterisk indicate a significant difference between the NIP and *OsRACK1B* overexpression lines (**, *P* < 0.01, Student’s *t*-test, n = 20).

### OsRACK1B plays a positive role in rice resistance against *Xoc*

To determine the subcellular localization of OsRACK1B, we co-expressed OsRACK1B-CFP with the nuclear marker ABF1-mCherry in rice protoplasts. The CFP signal was observed in the plasma membrane, cytoplasm, and nucleus (Figure 3c), indicating that OsRACK1B is a multi-compartmental protein. To further investigate the functional role of *OsRACK1B* in rice responses to *Xoc*, we generated stable transgenic rice lines constitutively overexpressing *OsRACK1B* under the control of the CaMV 35S promoter via *Agrobacterium tumefaciens*-mediated transformation. A total of fifteen independent T₀ transgenic lines were confirmed by PCR and subsequently advanced to the T₂ generation. Among these, two independent homozygous overexpression lines, designated RACK1BOE-2 and RACK1BOE-3, exhibited elevated OsRACK1B transcript abundance compared to wild-type NIP plants and were selected for subsequent functional analyses (Figure 3d). To evaluate the impact of *OsRACK1B* overexpression on BLS resistance, we inoculated transgenic and wild-type plants with *Xoc* strain GX01. At 14 days post-inoculation (dpi), both RACK1BOE lines displayed significantly reduced lesion lengths compared to wild-type controls, indicative of enhanced resistance to *Xoc* infection (Figure 3e,f).

We also generated *OsRACK1B* knockout mutants using CRISPR–Cas9-mediated genome editing. Two independent knockout lines, OsRACK1BKO-2 and OsRACK1BKO-8, were identified. PCR amplification and sequencing analysis revealed that OsRACK1BKO-2 carried a 2-bp deletion and a 1-bp insertion at the U6a-gRNA and U6b-gRNA target sites, respectively, whereas OsRACK1BKO-8 harbored a 1-bp insertion at both target sites (Figure 4a). Following GX01 inoculation, lesion lengths in the knockout plants were significantly greater than those in wild-type controls (Figure 4b,c). Together, these findings indicate that *OsRACK1B* functions as a positive regulator of resistance to BLS in rice.

**Figure 4.**
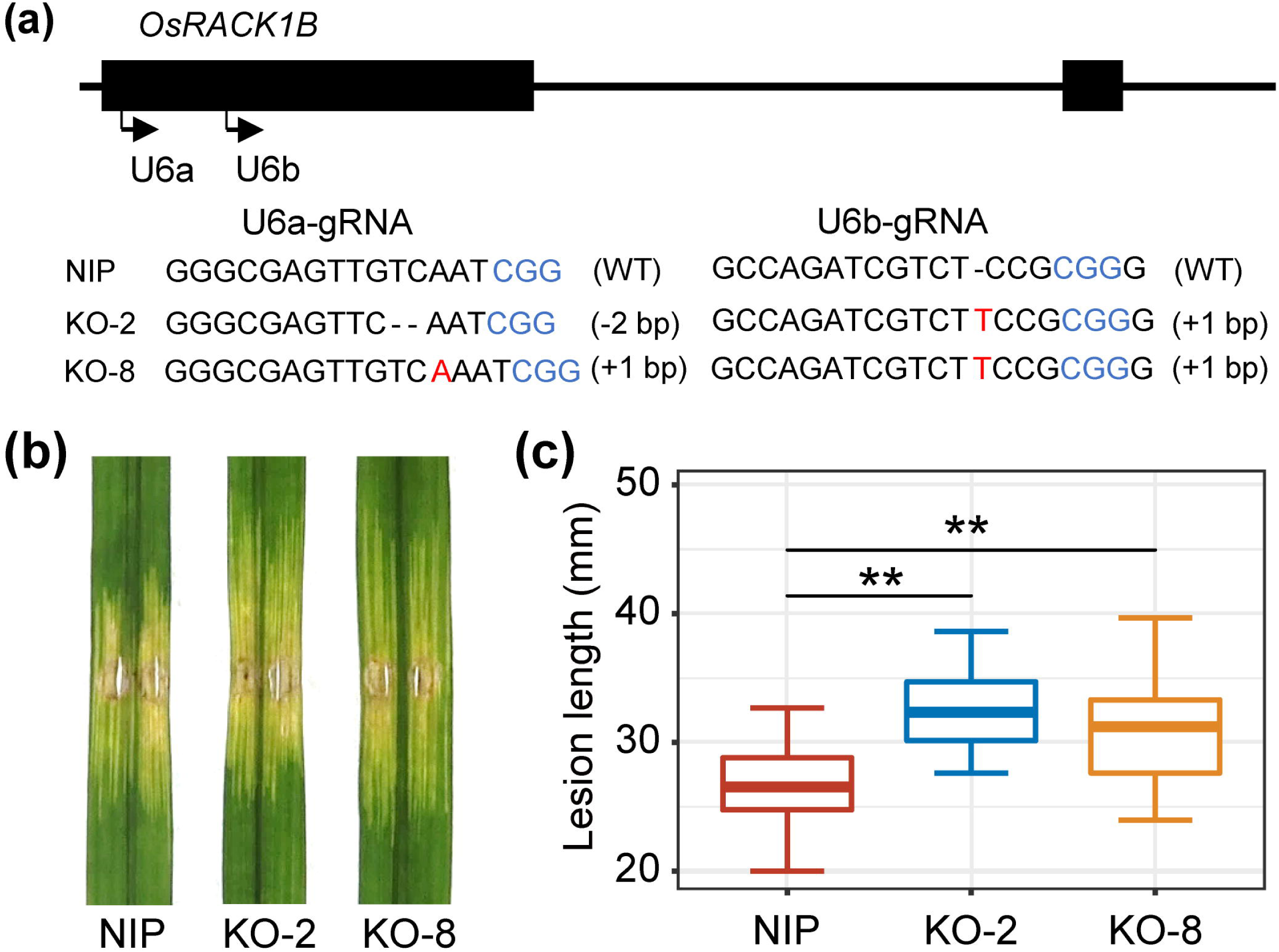
Loss of *OsRACK1B* function compromises rice resistance to *Xoc*. **(a)** Generation of *OsRACK1B* homozygous knockout mutants via CRISPR-Cas9 editing. The schematic illustrates the *OsRACK1B* gene structure and the targeted genomic locus. Blue nucleotides denote the protospacer adjacent motif (PAM); red letters indicate inserted bases; hyphens represent nucleotide deletions. **(b)** Phenotypes of lesion expansion in the NIP and *OsRACK1B* edited mutants at 14 days post-inoculation with GX01. **(c)** Lesion lengths in the NIP and *OsRACK1B* edited mutants at 14 days post-inoculation with GX01. Data are displayed as boxplots. Error bars represent maximum and minimum values. Center line, median; box limits, 25th and 75th percentiles. Asterisks indicate a significant difference between the NIP and OsRACK1B edited mutants (**, *P* < 0.01, Student’s *t*-test, n = 20).

### Knockout of *OsRACK1B* suppresses the rice immune response

To gain mechanistic insights into the role of OsRACK1B in rice innate immunity, we assessed callose deposition in *OsRACK1B* knockout mutants. Following inoculation with the Δ*hrcC* mutant strain, the knockout lines exhibited a marked reduction in callose accumulation in leaf tissues compared to wild-type plants (Figure 5a,b). In parallel, DAB staining indicated that Δ*hrcC*-induced ROS production was substantially attenuated in the knockout mutants relative to the wild type (Figure 5c). To further explore the impact of *OsRACK1B* on immune signaling, we quantified the transcript levels of two defense-related genes, *OsPBZ1* and *OsPAL1*, in the mutant lines. RT–qPCR analysis revealed a significant downregulation of both genes following Δ*hrcC* challenge in OsRACK1B knockout plants compared to wild-type controls (Figure 5d,e). These observations suggest that OsRACK1B contributes to PTI by regulating callose deposition, ROS accumulation, and defense gene expression.

**Figure 5.**
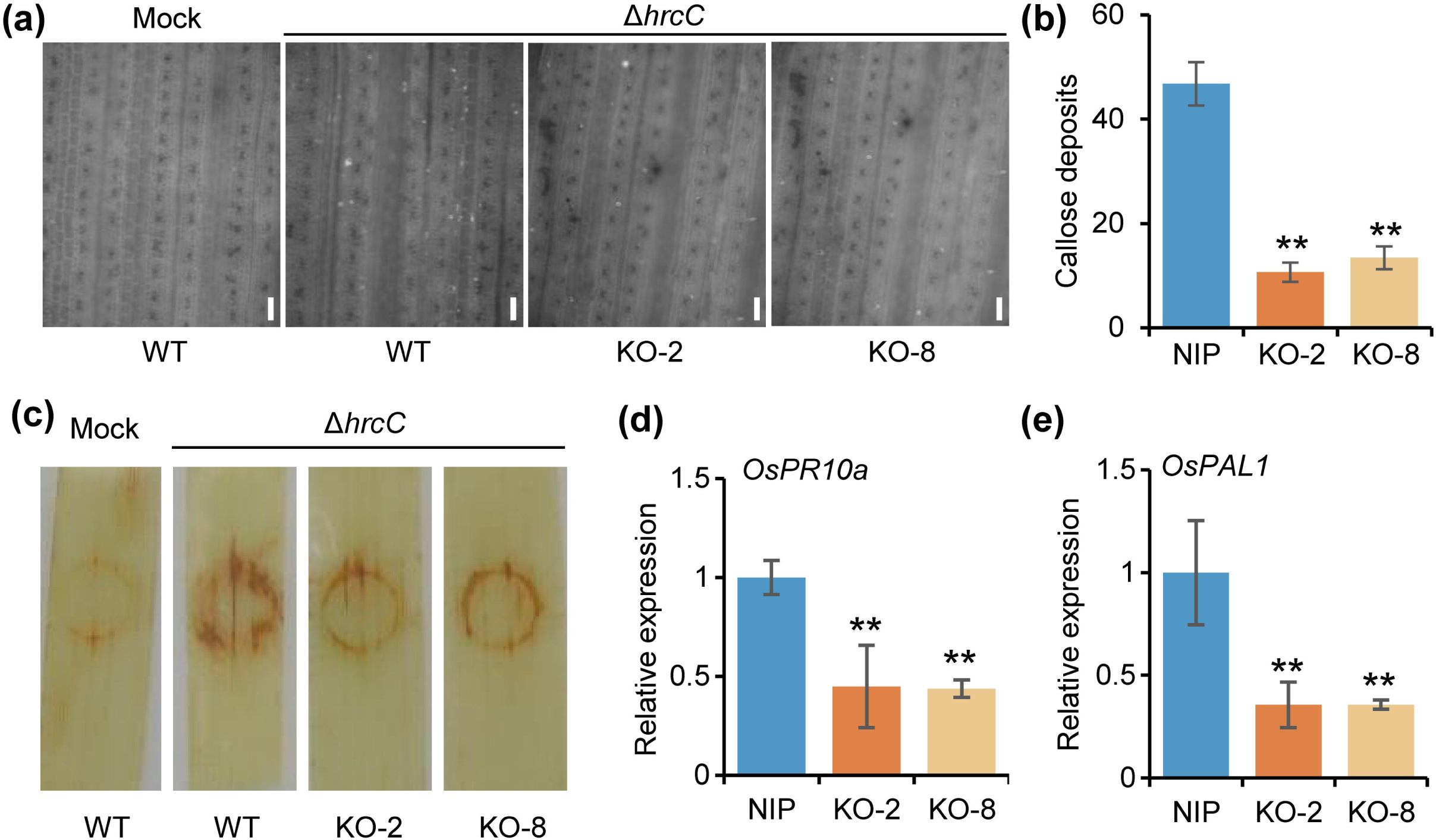
*OsRACK1B* is required for rice immune responses to *Xoc.* **(a)** Callose deposition in wild-type NIP and *OsRACK1B* knockout lines at 48 hours post-inoculation with Δ*hrcC*. Scale bar, 50 μm. **(b)** Quantification of callose deposits per leaf in NIP and OsRACK1B knockout plants. Data represent means ± standard deviations, n=5. Asterisks denote statistically significant differences between NIP and *OsRACK1B* knockout lines (**, *P* < 0.01, Student’s *t*-test). **(c)** DAB staining showing hydrogen peroxide (H₂O₂) accumulation in NIP and *OsRACK1B* knockout lines at 48 hours post-inoculation with Δ*hrcC*. **(d, e)** Relative transcript levels of *OsPBZ1* and *OsPAL1* in NIP and *OsRACK1B* knockout lines at 24 hours post-inoculation with Δ*hrcC*. Rice *actin* was used as an internal control. Data represent means ± standard deviations, n=3. Asterisks denote statistically significant differences between NIP and *OsRACK1B* knockout lines (**, *P* < 0.01, Student’s *t*-test).

### XopN impairs the OsRACK1B–OsRap2.6 interaction

The transcription factor OsRap2.6 plays a pivotal role in rice innate immunity through its interaction with RACK1 (Wamaitha et al., 2012; Yoshihisa et al., 2022). In the present study, we first confirmed the physical association between OsRap2.6 and OsRACK1B using Y2H assays (Figure S3). Given that the bacterial effector XopN also binds to OsRACK1B, we hypothesized that XopN might modulate the OsRACK1B–OsRap2.6 interaction. Therefore, we first employed AlphaFold3 prediction to investigate this hypothesis. The results showed that the predicted interaction score between OsRACK1B and OsRap2.6 was 0.65, which decreased substantially to 0.25 upon the addition of XopN, suggesting that XopN may inhibit the interaction between OsRACK1B and OsRap2.6 (Figure 6a). To validate this finding, we employed yeast three-hybrid (Y3H) assays to assess the OsRACK1B–OsRap2.6 interaction in the presence of either full-length. XopN or a truncated variant, XopN1–358. Notably, co-expression of either form of XopN significantly attenuated the interaction between OsRACK1B and OsRap2.6 (Figure 6b). These findings suggest that XopN competitively disrupts the association between OsRACK1B and OsRap2.6, potentially interfering with downstream immune signaling.

**Figure 6.**
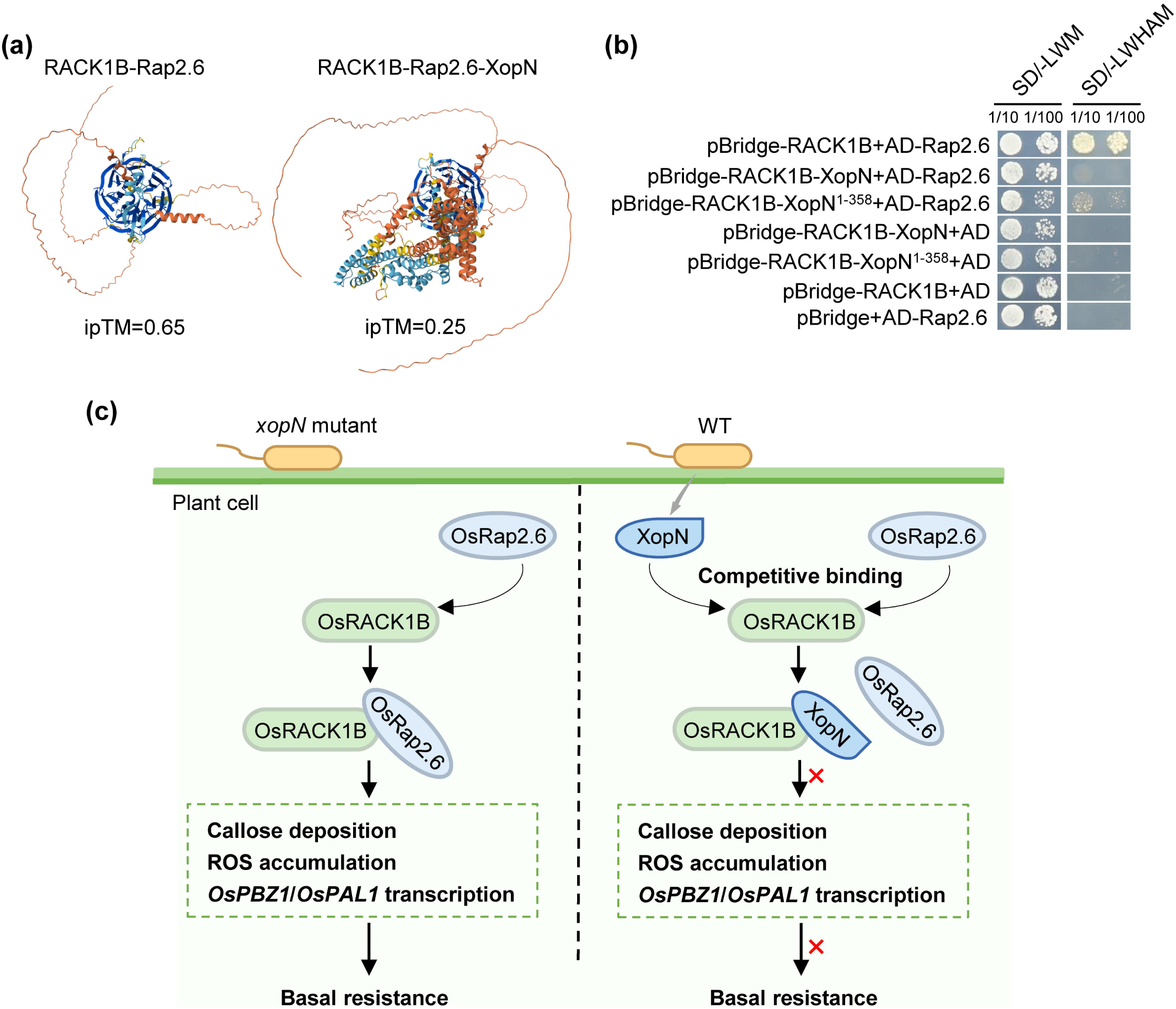
XopN competes with OsRap2.6 for binding to OsRACK1B. **(a)** AlphaFold predicts the effect of XopN on the OsRACK1B–Rap2.6 interaction. **(b)** Y3H assay was performed to evaluate the competitive binding dynamics between XopN and OsRap2.6 for OsRACK1B. Yeast strain Y2HGold was co-transformed with the indicated plasmid combinations and cultured on SD/-Leu-Trp-Met and the selective medium SD/-Leu-Trp-His-Ade-Met. Plates were incubated at 30 °C for 3 days prior to imaging. EV, empty vector control. SD/-LWM, SD/-Leu-Trp-Met; SD/-LWHAM, SD/-Leu-Trp-His-Ade-Met. **(c)** Schematic summary of XopN function in rice–*Xoc* interactions. During infection, the type III effector XopN is secreted by *Xoc* and translocated into rice cells, where it targets the scaffold protein OsRACK1B, a key component of rice immune signaling against *Xoc*. XopN binds to OsRACK1B in a manner that competitively interferes with the recruitment of the transcription factor OsRap2.6, thereby disrupting the formation of an OsRACK1B–OsRap2.6 immune complex. This disruption attenuates downstream immune responses, including callose deposition, ROS accumulation, and the transcriptional activation of defense-related genes, ultimately facilitating bacterial colonization of host tissue.

## Discussion

T3SEs are key virulence determinants in Gram-negative bacterial pathogens that actively manipulate host cellular processes to promote infection(Kay and Bonas, 2009). These effectors are known to specifically target host substrates, thereby suppressing immunity and enhancing pathogen proliferation. In the present study, we demonstrate that *Xoc* effector XopN facilitates bacterial infection by targeting the rice scaffold protein OsRACK1B, a positive regulator of innate immunity. During infection, XopN is translocated into host cells via the T3SS and competes with OsRap2.6 for binding to OsRACK1B. This competitive interaction leads to impaired immune signaling, as evidenced by reduced callose deposition, attenuated ROS accumulation, and downregulation of defense-related gene expression, thereby promoting successful colonization of host tissues by *Xoc* (Figure 6c).

To establish a successful infection, many host-adapted bacterial pathogens deploy T3SEs that suppress plant immune responses from within host cells. Previous research has demonstrated that *Xcv* effector XopN inhibits callose deposition in both tomato and *Arabidopsis thaliana* (Kim et al., 2009). In line with these findings, we observed that XopN from *Xoc* also suppresses callose accumulation in rice (Figure 1a,b), suggesting a conserved effector function across diverse plant species. Moreover, XopN inhibited ROS generation and downregulated the expression of defense-related genes (Figure 1c-e). Collectively, these data indicate that XopN enhances *Xoc* virulence by broadly attenuating multiple layers of basal immune defense.

Phytopathogen effectors have evolved to subvert host immunity through direct interactions with plant proteins, thereby undermining defense signaling pathways (Dodds and Rathjen, 2010). In the present work, we provide multiple lines of evidence by Y2H, in vitro pull-down and BIFC assays showing that XopN interacts with the rice scaffold protein OsRACK1B (Figure 2). Previous studies have shown that *Xcv* XopN is localized to both the PM and the cytoplasm (Kim et al., 2009). Our analysis of XopN localization in planta, using transient expression of XopN-GFP in rice protoplasts, supports this finding (Figure 1f). OsRACK1B is localized in the PM, cytoplasm, and nucleus (Supplementary Figure 3c), suggesting that the localization of OsRACK1B partially overlaps with that of XopN in plant cells. However, whether the interaction between XopN and OsRACK1B in these subcellular compartments is essential for full resistance remains unclear. Future studies will be necessary to delineate the precise subcellular locations of their interaction and further explore the molecular functions of XopN in plant immunity.

In *Arabidopsis*, the three RACK1 homologs AtRACK1A, AtRACK1B, and AtRACK1C have been implicated in a wide range of physiological processes, including developmental regulation, hormone signaling, innate immunity, and responses to abiotic stress (Chen et al., 2006; Cheng et al., 2015; Islas-Flores et al., 2015). Rice encodes two homologs, OsRACK1A and OsRACK1B. While overexpression of OsRACK1A has been reported to promote ROS accumulation and enhance resistance to *Ustilaginoidea virens* and *Magnaporthe oryzae* (Li et al., 2022; Nakashima et al., 2008), the functional role of OsRACK1B in immunity has remained largely unexplored. Here, we show that OsRACK1B expression is transcriptionally upregulated during the early stages of infection, as revealed by RT-qPCR analysis (Figure 3a-b), suggesting a role in basal defense against *Xoc*. This hypothesis was further substantiated by resistance assays in OsRACK1B-overexpression and knockout lines (Figure 3 c-e and Figure 4). Further functional analysis revealed that OsRACK1B is essential for Δ*hrcC*-triggered callose deposition, ROS production, and transcriptional activation of defense-related genes in rice (Figure 5). Collectively, these findings identify OsRACK1B as a positive regulator of rice immunity and suggest that it may serve as a critical node in the defense signaling network against *Xoc* infection.

Scaffold protein RACK1 serves as a pivotal integrative hub in eukaryotes, coordinating diverse signaling pathways through interactions with numerous protein partners (Fu et al., 2024; Islas-Flores et al., 2015). Prior studies have demonstrated that RACK1 forms a complex with the transcription factor Rap2.6, a module implicated in rice immunity against *M. oryzae* (Guo and Sun, 2017; Wamaitha et al., 2012). More recently, OsRap2.6 has been shown to regulate NLR Xa1-mediated immunity in response to TAL effector recognition, underscoring its central role in pathogen resistance signaling (Yoshihisa et al., 2022). Here, we report that the *Xoc* effector XopN disrupts the interaction between OsRACK1B and OsRap2.6 (Figure 6a,b). However, the functional consequence of this disruption on OsRap2.6 remains unclear. Based on structural and functional considerations, we speculate that XopN may affect the stability or transcriptional activity of OsRap2.6. This mechanism is reminiscent of the interaction between *Xcv* effector XopN and the tomato proteins TFT1 and TARK1, where XopN promotes the formation of TFT1-TARK1 complex (Kim et al., 2009; Taylor et al., 2012). Notably, in contrast to the stabilizing effect observed in tomato, XopN in rice appears to dismantle the OsRACK1B–OsRap2.6 complex, suggesting a species-specific modulation by this evolutionarily conserved effector. Structural elucidation of XopN would likely reveal the molecular basis of its interaction with the OsRACK1B–OsRap2.6 subcomplex and offer mechanistic insights into its mode of action.

In conclusion, our findings reveal a potential pathogenic mechanism through which *Xoc* type III effector XopN targets the rice scaffold protein OsRACK1B to attenuate basal immunity and promote bacterial colonization. This work deepens current understanding of non-TALE-mediated host–pathogen interactions and holds implications for the development of disease-resistant rice cultivars.

## Materials and Methods

### Plants and bacterial strains

Rice cultivar Nipponbare was used as the wild-type (WT) control in this study. For the overexpression of OsRACK1B, its coding sequence was PCR-amplified and cloned into the pCAMBIA1301-HA binary vector. To generate knockout lines, two guide RNAs targeting the coding region of OsRACK1B were designed using the CRISPR-GE online toolkit (Xie et al., 2017). The gRNA expression cassettes were subsequently inserted into the pYLCRISPR/Cas9 binary vector. Both overexpression and CRISPR constructs were introduced into *Nipponbare* via *A. tumefaciens*-mediated transformation (Wuhan BioRun Biosciences Co., Ltd.). Rice plants were cultivated in greenhouse conditions under a 14-hour light (30°C)/10-hour dark (28°C) photoperiod.

*E. coli* and *A. tumefaciens* strains were propagated in LB medium at 37°C and 28 °C, respectively. *Xoc* strains were cultured at 28 °C in NB medium (polypeptone 5g/L, beef paste 3g/L, yeast extract 1g/L, sucrose 10g/L). Virulence assays were conducted as previously described (Wang et al., 2024). Briefly, *Xoc* strains were resuspended in sterile distilled water to an optical density of OD_600_ = 0.5 and infiltrated into rice leaves using needleless syringes. Lesion lengths were recorded, and photographs were taken 14 days post-inoculation.

### RNA analysis

Total RNA was extracted from rice leaf tissue using TRIzol Reagent (Invitrogen, Cat# 15596018, USA) according to the manufacturer’s protocol. First-strand cDNA synthesis was performed with the RevertAid First Strand cDNA Synthesis Kit (Thermo Fisher Scientific, Cat# K1622, USA). Quantitative real-time PCR (RT-qPCR) was conducted using the qTOWER 2.2 Real-Time PCR Detection System (Analytik Jena, Germany) with ChamQ SYBR Color qPCR Master Mix (Vazyme, Cat# Q411-02, China), following the manufacturer’s instructions. OsACTIN1 was used as the internal reference gene to normalize transcript levels. Relative gene expression was calculated using the 2^-ΔΔCT^ method. Primer sequences used in this study are listed in Table S1.

### Callose deposition and DAB staining

Leaves from 2–3-week-old rice seedlings inoculated with *Xoc* were fixed in ethanol: acetic acid (3:1, v/v) for 5 h, with frequent replacement of the solution until fully decolorized. Samples were subsequently rehydrated through a graded ethanol series (70% for 2 h, 50% for 2 h), followed by incubation in distilled water overnight. After triple rinsing with water, the tissues were cleared by incubation in 10% (w/v) NaOH for 1 h. Leaves were then washed four times with water and stained in 0.01% aniline blue solution (150 mM K₂HPO₄, pH 9.5) for 4 h. Callose deposits were visualized immediately using a fluorescence microscope equipped with a UV filter(Deb et al., 2020).

For DAB staining, inoculated rice leaves were collected and incubated in DAB solution (1 mg/mL, Sigma Aldrich) for 24 h with gentle agitation. Leaves were subsequently cleared in a decolorizing solution composed of ethanol, acetic acid, and glycerol (3:1:1, v/v/v) before imaging(Deb et al., 2020).

### Y2H assay and Y3H assay

The Matchmaker gold Y2H system (Clontech) was used to screen XopN-interacting proteins. The XopN coding sequence was cloned into the bait vector pGBKT7 and then transformed into the yeast strain Y2H gold. A rice cDNA library was constructed into the prey vector pGADT7 using mRNA extracted from rice leaves according to the manufacturer’s protocol. Positive interactors were initially selected on SD/-Leu/-Trp/-His/-Ade dropout medium. To validate the interaction between XopN and OsRACK1B, their coding sequences were individually cloned into pGBKT7 and pGADT7, respectively, and co-transformed into the Y2HGold strain. Transformed yeast colonies were selected on SD/-Leu/-Trp plates, and interactions were assessed by growth on quadruple dropout medium (SD/-Leu/-Trp/-His/-Ade).

For the Y3H assay, the pBridge vector was used to express both XopN and OsRACK1B. The pGADT7 vector was used to express OsRap2.6. The recombinant constructs were co-transformed into the yeast strain Y2HGold. Transformants were selected on SD/-Met/-Trp/-Leu medium, and protein – protein interactions were assessed by growth on SD/-Met/-Trp/-Leu/-His/-Ade selective plates.

### Pull-down assay

The XopN coding sequence was cloned into the vector pET28a to generate His-XopN fusion protein. The coding sequence of OsRACK1B was cloned into the vector pGEX-6p-1 to generate GST-OsRACK1B fusion protein. His-XopN and GST-OsRACK1B were expressed in *E. coli* strain BL21 and were purified with Ni-NTA resin and GST resin, respectively (Transgen). His-XopN proteins were immobilized on Ni-NTA resin and incubated with GST-OsRACK1B or GST at 4°C for 4 h with gentle shaking. After five times washing with 1 × PBS (4.2 mM Na_2_HPO_4_, 2 mM KH_2_PO_4_, 140 mM NaCl, and 10 mM KCl), the mixtures were eluted using the elution buffer. Last, the elution samples were boiled with 5 × SDS loading buffer at 100°C for 5 min for immunoblot analysis.

### BIFC assay

The pXY104 and pXY106 plasmids were used for the BIFC assay(Fu et al., 2025). The coding sequence of XopN was cloned into pXY106 to make the XopN-nYFP fusion protein. The OsRACK1B coding sequence was cloned into pXY104 to make the OsRACK1B-cYFP fusion protein. The constructs were co-expressed in rice protoplasts. The YFP fluorescence was observed using a confocal laser scanning microscope (Leica TCS SP8 MP).

### Subcellular localization

To examine the subcellular location of XopN and OsRACK1B, full-length cDNAs were constructed into vector pBWA(V)HS-GFP and pBWA(V)HS-GFP to make the fusion proteins XopN-GFP and OsRACK1B-CF, respectively. The recombinant plasmids were transfected into rice protoplasts, and the fluorescence was observed using a confocal laser scanning microscope at 24 h after transfection.

### Sequence alignment and phylogenetic analysis

Orthologs of XopN from various bacterial species were identified and retrieved from the NCBI database. Protein sequences were aligned using the Constraint-based Multiple Alignment Tool (COBALT), and the resulting alignments were visualized with the online software ESPript 3.0 (Robert and Gouet, 2014). NCBI BLASTP was used to select and download the orthologs of OsRACK1B in other plants. A phylogenetic tree was constructed with MEGA 7.0 using the maximum likelihood (ML) method and 1,000 bootstrap replicates.

### Accession numbers

The sequence data from this article can be found on the Kyoto Encyclopedia of Genes and Genomes website (https://www.kegg.jp/) under the following numbers: *xopN* (*Xoc_0350*), *OsRACK1B* (*Os05g0552300*), *OsPBZ1* (*Os12g0555500*), *OsPAL1* (*Os02g0626100*), *OsRap2.6* (*Os04g0398000*) and *OsACTIN* (*Os03g0718100*).

## Author contributions

S.H. conceived the research and designed the experiments. J.W., Z.N., X.C., Y.Z. and H.Q. performed the experiments; J.W., Z.N. and H.W. analysed the data. J.W. wrote the manuscript. S.H., J.T., and Y.H. revised the manuscript. All authors have read and approved the final manuscript.

## Acknowledgments

This research was supported by the National Natural Science Foundation of China (31860032 and 32360045).

## Data availability statement

All relevant data can be found within the manuscript and its supporting materials.

## Supporting Information

Additional Supporting Information may be found online in the Supporting Information section at the end of the article.

**Figure S1.** Sequence alignment of XopN and its homologs.

**Figure S2.** Sequence analysis of OsRACK1B.

**Figure S3.** Validation of the interaction between OsRACK1B and OsRap2.6.

**Table S1** Primers used in this study.

