## Supplemental file for "*Xanthomonas* type III effector XopN targets scaffold protein OsRACK1B to suppress rice immunity"

### Supporting Information

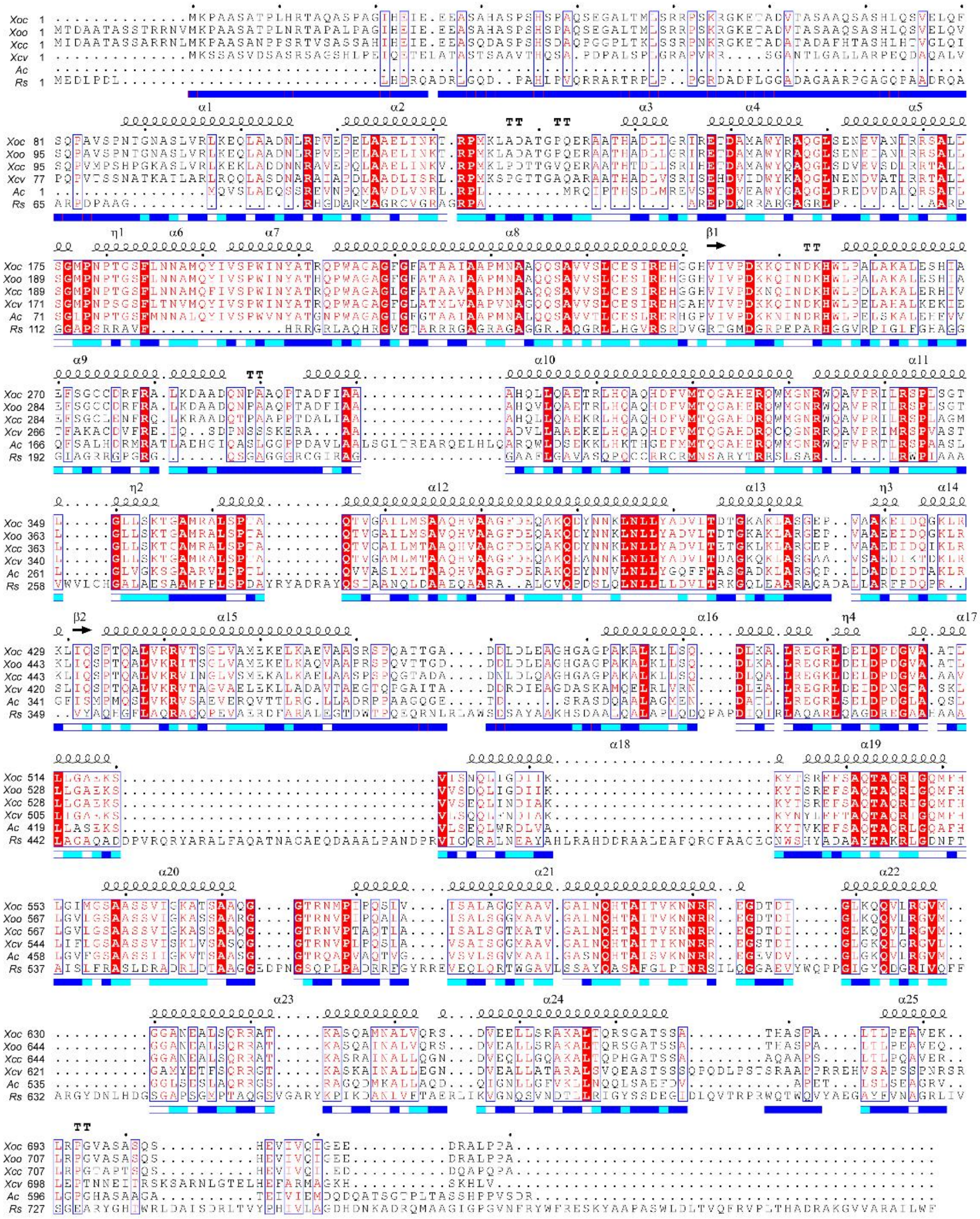

**Figure S1. Sequence alignment of XopN and its homologs.** XopN homologs were identified by BLAST searches against full-length protein sequence of XopN. The protein sequences of XopN and its homologs from *X. oryzae pv. oryzae* (*Xoo*), *X. campestris* pv. *vesicatoria* (*Xcv*), *Acidovorax citrulli* (*Ac*) and *Ralstonia solanacearum* (*Rs*), were aligned with COBALT, and were then were visualized with the online software ESPript 3.0. The secondary structures of the OsRACK1B are shown above the alignment: α (α-helix), arrows (β-strands), TT (β-turns), and η (310-helix). Identical and similar residues are boxed in blue.

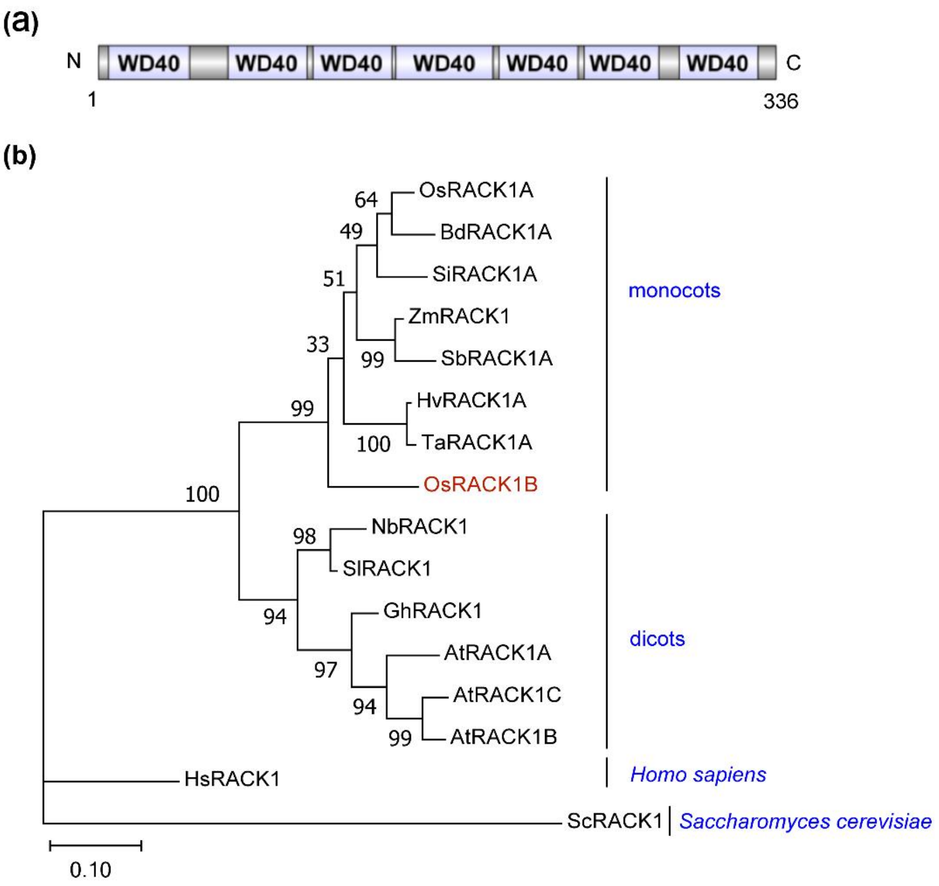


**Figure S2. Sequence analysis of OsRACK1B.** (A) The domain architecture of OsRACK1B was predicted using the SMART database. (B) Phylogenetic analysis of OsRACK1B and its homologous proteins. The phylogenetic tree was constructed using MEGA 7.0 software based on the maximum likelihood method, with bootstrap values derived from 1,000 replications. The proteins are from Oryza sativa (Os), Brachypodium distachyon (Bd), Setaria italica (Si), Zea mays (Zm), Sorghum bicolor (Sb), Hordeum vulgare subsp. Vulgare (Hv), Triticum aestivum (Ta), Nicotiana benthamiana (Nb), Solanum lycopersicum (Sl), Gossypium hirsutum (Gh), Arabidopsis thaliana (At), Homo sapiens (Hs) and Saccharomyces cerevisiae (Sc).

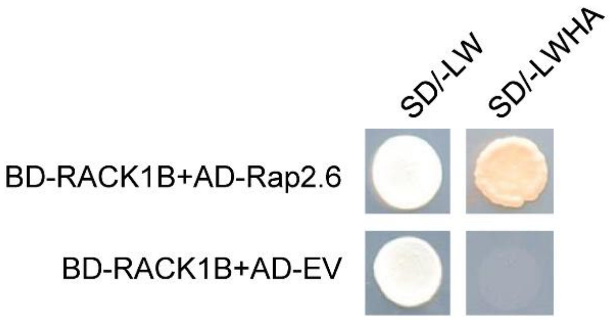


**Figure S3. Validation of the interaction between OsRACK1B and OsRap2.6.** Yeast strain Y2HGold was co-transformed with the indicated plasmid combinations and cultured on SD/-Leu-Trp and the selective medium SD/-Leu-Trp-His-Ade. Plates were incubated at 30 °C for 3 days prior to imaging. SD/-LW, SD/-Leu-Trp; SD/-LWHA, SD/-Leu-Trp-His-Ade.

**Table S1** Primers used in this study.

| **Primer** | **Sequence (5' to 3')** | **description** |
|---|---|---|
| OsACTIN-qRT-F | GAGACCTTCAACACCCCTGCT | qRT-PCR |
| OsACTIN-qRT-R | TCACCAGAGTCCAACACAATACCT | qRT-PCR |
| OsRACK1B-qRT-F | GCCAGGAGTCGCTCACCC | qRT-PCR |
| OsRACK1B-qRT-R | GGTGGACGGGTTGGTGATG | qRT-PCR |
| OsPBZ1-qRT-F | ATCGTGGATGGCTACTATGGC | qRT-PCR |
| OsPBZ1-qRT-R | GACATTTCTGCGGCTCTCATT | qRT-PCR |
| OsPAL1-qRT-F | CGCCATGGCCTCCTACTG | qRT-PCR |
| OsPAL1-qRT-R | GCTCTGGACATGGTTGGTGAT | qRT-PCR |
| AD-XopN1-358-F | ATGGAGGCCAGTGAATTCATGAAACCTGCTGCATCTGC | Y2H |
| AD-XopN1-358-R | TCGATGCCCACCCGGGTGTTACATTGCGCCGGTCTTG | Y2H |
| AD-XopN353-720-F | ATGGAGGCCAGTGAATTCAGCAAGACCGGCGCAAT | Y2H |
| AD-XopN353-720-R | ATGGAGGCCAGTGAATTCAGCAAGACCGGCGCAAT | Y2H |
| AD-XopN1-187-R | TCGATGCCCACCCGGGTGTTAATTGTTGAGGAAGCTGCCGGT | Y2H |
| AD-XopN182-305-R | TCGATGCCCACCCGGGTGTTACAGCAATTGATGAGCCGC | Y2H |
| AD-XopN182-35 | ATGGAGGCCAGTGAATTCGCTCATCAATTG | Y2H |

| | | |
|---|---|---|
| 8-R | CTG | |
| AD-XopN240-358-F | ATGGAGGCCAGTGAATTCGGGCACGTCATCGTGCC | Y2H |
| AD-XopN300-358-F | ATGGAGGCCAGTGAATTCGCTCATCAATTGCTGCAAG | Y2H |
| BD-OsRACK1B WD1-6-F | GCCATGGAGGCCGAATTCATGGCGGGCCAGGAGT | Y2H |
| BD-OsRACK1B WD1-6-R | AGGTCGACGGATCCCCGGCTAGTCCTGCATGACGAGCTT | Y2H |
| BD-OsRACK1B WD2-7-F | GCCATGGAGGCCGAATTCATCACCAACCCGTCCAC | Y2H |
| BD-OsRACK1B WD2-7-R | AGGTCGACGGATCCCCGGCTAGATTGCATAGCCGCC | Y2H |
| BD-OsRACK1B WD7-F | GCCATGGAGGCCGAATTCCTCAAGCCAGAAGTCCAGGCCT | Y2H |
| nYFP-XopN-F | GGGTCTAGAATGAAACCTGCTGCATCTGCC | BIFC |
| nYFP-XopN-R | GGGGTCGACCGCCGGCGGCAGTGCCC | BIFC |
| RACK1B-cYFP-F | GGGTCTAGAATGGGGGCCAGGAGTCGCT | BIFC |
| RACK1B-cYFP-R | TTTGTCGACCTAGATTGCATAGCCGCCAA | BIFC |
| XopN-GFP-F | GGGTCTAGAATGAAACCTGCTGCATCTGCC | Subcellular localization |
| XopN-GFP-R | GGGGGTACCCGCCGGCGGCAGTGCCC | Subcellular localization |
| RACK1B-CFP-F | GGGTCTAGAATGGCGGGCCAGG AGTCGCT | Subcellular localization |
| RACK1B-CFP-R | GGGGGTACCGATTGCATAGCCGCCAAACC | Subcellular localization |
| His-XopN-F | GGAAGATCTATGAAACCTGCTGCATCTGCC | Pull down |
| His-XopN-R | CCGCTCGAGCGCCGGCGGCAGTGCCC | Pull down |
| GST-RACK1B-F | CCGGAATTCATGGCGGGCCAGGAGTCGCT | Pull down |
| GST-RACK1B-R | GGGGTCGACGATTGCATAGCCGCCAAACC | Pull down |
| pBridge-OsRACK1B-F | ACTGTATCGCCGGAATTCATGGCGGGCCAGGAGT | Y3H |
| pBridge-OsRACK1B-R | AGGTCGACGGATCCCCGGCTAGATTGCATAGCCGCC | Y3H |
| pBridge-OsRACK1B-XopN-F | AGAAAGGTGGCGGCCGCAATGAAACCTGCTGCATCTGC | Y3H |
| pBridge-OsRAC | CGAAGATCTTCGGGCTAATTACGCCGGCG | Y3H |

| | | |
|---|---|---|
| K1B-XopN-R | GCAGT | |
| pBridge-OsRACK1B-XopN1-358-R | CGAAGATCTTCGGGCTAATTACATTGCGCCGGTCTTG | Y3H |
| U-F | CTCCGTTTTACCTGTGGAATCG | Gene editing rice construction |
| gR-R | CGGAGGAAAATTCCATCCAC | Gene editing rice construction |
| Pps-R | TTCAGAGGTCTCTACCGACTAGTCACGCGTATGGAATCGGCAGCAAA | Gene editing rice construction |
| Pgs-2 | AGCGTGGGTCTCGTCAGGGTCCATCCACTCCAAGCTC | Gene editing rice construction |
| Pps-2 | TTCAGAGGTCTCTCTGACACTGGAATCGGCAGCAAAGG | Gene editing rice construction |
| Pgs-L | AGCGTGGGTCTCGCTCGACGCGTATCCATCCACTCCAAGC | Gene editing rice construction |
| OsRACK1B-gRT1 | ATGAAGGGCGAGTTGTCAATGTTTTAGAGCTAGAAAT | Gene editing rice construction |
| OsRACK1B-U6aT1 | ATTGACAACTCGCCCTTCATCGGCAGCCAAGCCAGCA | Gene editing rice construction |
| OsRACK1B-gRT2 | CGGAGACGATCTGGCGGTTGCAACACAAGCGGCAGC | Gene editing rice construction |
| OsRACK1B-OsU6bT2 | CGGAGACGATCTGGCGGTTGCAACACAAGCGGCAGC | Gene editing rice construction |
| OsRACK1B-KO-VF | TCTCCTAGCCTTCCCCTCCT | CRISPR/Cas9 mutant identification |
| OsRACK1B-KO-VR | GGCTTGAGGTCCTGCATGAC | CRISPR/Cas9 mutant identification |
| OsRACK1B-OE-F | GGGCCCGGTACCGAGCTCATGGCGGGCCAGGAGTC | Overexpression vector construction |
| OsRACK1B-OE-R | GCGGCCGCCACCGCGGTGGATTGCATAGCCGCCAAAC | Overexpression vector construction |
| OsRACK1B-OE-VF | ACTCCCTCTGCTTCTCGCC | PCR Identification for overexpression |
| OsRACK1B-OE-VR | CTGCAGCCCCTAAGCGC | PCR Identification for overexpression |
